# An Integrative Multitiered Computational Analysis for Better Understanding the Structure and Function of 85 Miniproteins

**DOI:** 10.1101/2025.01.31.635936

**Authors:** Reethika Veluri, Gareth Pollin, Jessica B. Wagenknecht, Raul Urrutia, Michael T. Zimmermann

**Author notes:** Corresponding Author: Michael T. Zimmermann.

## Abstract

**Background:** Miniproteins, defined as polypeptides containing fewer than 50 amino acids, have recently elicited significant interest due to an emerging understanding of their diverse roles in fundamental biological processes. In addition, miniprotein dysregulation underlies human diseases and is a significant focus for biotechnology and drug development. Notably, the human genome project revealed the existence of many novel miniproteins, most of which remain uncharacterized. This study reports an approach for analyzing and scoring previously uncharacterized miniproteins by integrating knowledge from classic sequence-based bioinformatics, computational biophysics, and system biology annotations. Our results demonstrate that these approaches provide novel information on the structure-function relationship of these molecules with a particular focus on their biomedical relevance.

**Methods:** We identified 85 human miniproteins using a simple multi-tier approach. First, we performed a sequence-based analysis of these proteins using several algorithms to identify regions of structural and functional importance. Protein-protein interactions and gene ontology annotations were used to analyze miniprotein function. Then, we predicted miniprotein three-dimensional structures using AI-based methods and peptide modeling to determine their relative yields for these understudied polymers. Subsequently, we used several computational biophysics methods and structure-based calculations to annotate and evaluate results from both algorithms.

**Results:** We find several relations between predicted structure and functional properties to assign these proteins to several groups with similar properties. Sequence-based analysis leads us to identify motifs and residues that link structure-to-function for most of these proteins. We suggest novel miniprotein functions, such as thymosin beta proteins regulating the shelterin complex through TERF1 and POT1 interactions, FAM86JP and FAM66E participating in endocytic processes, and BAGE1 influencing chromatin remodeling through interaction with nuclear proteins. Further, known functions of miniproteins, such as STRIT1, STMP1, and SLN, were supported. Finally, structure-based scoring led us to build 3D models that provided complementary information to ontologies. We identify that structural propensity is not strictly dependent on polymer length. In fact, in this dataset, peptide-based algorithms may have advantages over AI-based algorithms for certain groups of miniproteins.

**Conclusion:** This analytic approach and resulting identification and annotation of miniproteins adds much to what is currently known about miniproteins. Our determination of novel properties of miniproteins bears significant mechanistic and biomedical relevance. We propose novel functions of miniproteins, which expands our understanding of their potential roles in cellular processes. And, we practically identify which sequence and structure-based tools provide the most information, aiding future studies of miniproteins.

## Introduction

The human cell can be described as an intricate system of biological processes with various types and functions. With advancements in scientific tools and methods, we have increased our understanding of these biological processes and their significant players. Gene products such as proteins remain at the forefront of physiology. However, due to technical aspects of short-read sequencing, a biological blind spot has emerged. Little is known about miniproteins, typically defined as polypeptides under 50-80 amino acids (CrookNairn & Olson 2020) (up to 10 kDa), although there is no strict consensus for their definition. While miniproteins present technical challenges to study, previous studies describe the potential that miniproteins have in genomic sciences. The miniprotein spectrum, including from other species, is known to contain signaling peptides, hormones, toxins, and protease inhibitors (Wang et al. 2008). Due to their short length, they are often assumed to be intrinsically disordered, yet there are experimental examples of their adoption of stable structures (Anderson et al. 2015; Hobbs et al. 2012; OjedaWang & Craik 2016; Pueyo & Couso 2008). Previous experiments on miniproteins have also shown potential function as scaffolding proteins (ZollerHaberkorn & Mier 2011) and pharmacological application as both inhibitors and activators of GPCRs (CrookNairn & Olson 2020). However, it must be noted that most miniproteins have relatively unknown or uncharacterized functions.

In previous genome-wide studies, data corresponding to miniproteins has often been filtered out due to low mappability in the genome, low conservation, and small size (Tripathi et al. 2021). We typically assume that the more conserved a genomic region is, the more functionally important it is. However, in some instances, that paradigm misses the unique regions and evolutionary changes that define our human species. Additionally, small nucleotide fragments are removed from library preparations, leading to technical features that put miniproteins at a disadvantage in genomics research. Additionally, miniproteins that have known roles in human diseases may not have a known role in normal physiology, and vice versa. This raises questions over how much of the normal function and physiology is known on miniproteins and if they have unrecognized contributions to additional diseases. By filling this blind spot, miniprotein research has the potential to advance biological knowledge, providing opportunities for more effective and precise treatment of disease. However, with data on miniproteins being few and far between, understanding this function is difficult and relies on integrating various sources of information.

In this report, we determine a process for analyzing and scoring critical features that serve as structural-functional signatures of previously uncharacterized miniproteins using sequence-based bioinformatics with extensive system biology annotations and paired with three-dimensional structure prediction. We intend to classify miniproteins further and infer possible biological function(s) using protein structure annotations and scores. Combined, these approaches provide novel information describing the structure-function relationship of miniproteins with potential biomedical relevance. Combining information across multiple sequence-based and structure-based algorithms provided the most current data about potential functions. Thus, both the analytical approach and the resulting new knowledge should be considered to extend the characterization of the extensive repertoire of miniproteins that remain to be characterized.

## Materials & Methods

### Process for Miniprotein Identification and Annotation

Human proteins under 50 amino acids were obtained from UniProt (UniProt 2019), a comprehensive resource for protein sequences and annotation data. The preliminary list comprised proteins with the Swiss-Prot Reviewed status, indicating that the protein had been experimentally verified or computationally predicted. Subsequently, the acquired sequences underwent analysis through protein annotation and sequence scoring web tools, facilitating miniprotein classification and function analysis. For Gene Ontology (GO) information, the web tool MyGene.info (Lelong et al. 2022; WuMacleod & Su 2013; Xin et al. 2016) was utilized. Additionally, InterPro (Paysan-Lafosse et al. 2022), neXtProt (Gaudet et al. 2017), CATH (Sillitoe et al. 2021), and The Eukaryotic Linear Motif resource (Kumar et al. 2019) were employed for their information on domains, motifs, and other associated annotations. IUPred (MészárosErdős & Dosztányi 2018) was integrated for its predictions on the intrinsic disorder. To further visualize miniprotein categorization, Multiple Sequence Alignment was performed on the acquired dataset using the MAFFT algorithm in Jalview2 (Waterhouse et al. 2009).

### Sequence-Based Analysis

Post-translational modifications for Miniproteins were retrieved from the Phosphosite Plus database (v6.7.1.1)(Hornbeck et al. 2015). All post-translational modifications (PTMs) present in these miniproteins were systematically documented. DOG software (v2.0)(Ren et al. 2009) was employed to create schematic diagrams annotating the positions of PTMs and any associated domain structures. To assess the potential functional and structural significance of PTMs, a conservation analysis was performed using ConSurf (Ashkenazy et al.) for the Humanin and Thymosin families.

### Miniprotein Predicted Interactome

To construct a comprehensive interactome for miniproteins, we utilized various protein interaction databases, including Biogrid (v4.4.229)(Oughtred et al. 2021), HIPPIE (v2.3)(Alanis-LobatoAndrade-Navarro & Schaefer 2016), Human Reference Interactome Mapping Project (Luck et al. 2020), IntAct (1.0.4)(Orchard et al. 2013), and IID (v2021-05)(Kotlyar et al. 2016). This approach incorporated both high-throughput datasets and predictive interactions based on sequence analysis. The resulting protein-protein interaction network was visualized using Cytoscape (v3.9.1)(Shannon et al. 2003). For ontological analysis, Enrichr (Xie et al. 2021) was employed to generate gene ontology profiles. Subsequently, these profiles were visualized as a dot blot using RStudio (v4.1.1) with the clusterProfiler package (v4.0)(Yu et al. 2012).

### Structure-Based Analysis

Utilizing the miniprotein sequences, two differing structure prediction tools were applied to determine the possible fold of proteins – while integrating the previously acquired sequence annotations. This involved contrasting the commonly used AI deep learning tools ALPHAFOLD2 (Jumper et al. 2021) and PEPFOLD3 (Lamiable et al. 2016), a de novo approach in peptide structure predictions. AlphaFold2 provided five structures per miniprotein with at least eight amino acids in length. Each of the structures was ranked by their pLDDT score (predicted local distance difference test score), representing the percentage of interatomic distance between side chain angles, thus giving local structure confidence. PEPFOLD3 provided ten models per miniprotein with at least five amino acids. PEPFOLD3 models were ranked by their sOPEP score, which is based on the energies of a structure using coarse-grained representation. Extracting scores from the given structures allowed for comparing the tools’ performance in terms of confidence and accuracy. Following score compilation, we compared the structures’ TM scores (provided by PEPFOLD3 and obtained from AlphaFold using the TM-score calculator developed by the Zhang Lab (Zhang & Skolnick 2004). Qmean scores were also compared.

Miniprotein 3D structures were further analyzed using PyMOL. This two-fold analysis involved intraprotein analysis, which compares alternative models of the same miniprotein, and interprotein analysis, which compares the top model of miniproteins within the same gene family. This was done for five miniprotein-rich families identified in our dataset: BAGE proteins, Humanin proteins, Keratin-associated proteins, Thymosin Beta proteins, and T-cell receptor proteins. Using the best model for each protein (determined by AlphaFold2’s calculated pLDDT score), RMSDs between all proteins within the same family were calculated using the CE algorithm implemented in PyMOL and recorded in a square matrix.

To comprehend the broader scope of the miniprotein dataset, structure matching was conducted using the Dali server (Holm 2022). All five models per miniprotein (obtained during structure prediction) were entered into Dali, and the top five matched models per result were chosen based on Z-scores. The CE algorithm was again utilized to visualize the structure matches, seeking possible patterns and resemblances.

## Results

### Identification of Miniproteins, Their Structural Features, Functional Annotations, and Disorder Predictions

Using our selection criteria (see Methods), we generated a dataset of 85 miniproteins containing between 2 and 49 amino acids (**Fig. 1A**). The distribution of miniprotein size shows the potential for the dataset to contain protein families, with clusters representing sequences between 15 and 25 amino acids and a group of related sequences around 45 amino acids in length. Several tools were used to obtain a preliminary understanding of these polymers and examine possible functional subgroups. Several miniprotein sequences are shorter than is required for some annotation tools, representing another practical challenge for exploring miniprotein function. For example, CATH, a popular protein topologic database, had no information available for any member of the miniprotein dataset. Yet, other protein annotation tools provide insight into the structure and function of miniproteins. For example, the Gene Ontology (GO) (Ashburner et al. 2000; Consortium et al. 2023) returned information for 73 of the 85 miniproteins at a gene level. The quantity of GO terms varied; of the 73 miniproteins with GO results, 53 had Biological Process GO terms, 61 had Cellular Component GO terms, and only 21 had Molecular Function GO terms. Some miniproteins had multiple GO terms per category, typically for larger, more historically researched miniproteins. Yet, another annotation tool, InterPro (Paysan-Lafosse et al. 2022), annotated 46 out of the 85 miniproteins with protein family and domain information, partly because proteins needed to be longer than 17aa. Amongst these annotations, 13 families and superfamilies were identified (**Table 1**), as well as two domains (MKRN1 C-terminal Domain and Domain of unknown function 4535), 11 disordered proteins, 12 signaling peptides, and five transmembrane proteins. We identified patterns among miniprotein families (notably the Humanin protein family), domains, and superfamilies. Therefore, these understudied molecules likely have shared functional relationships.

**Figure 1:**
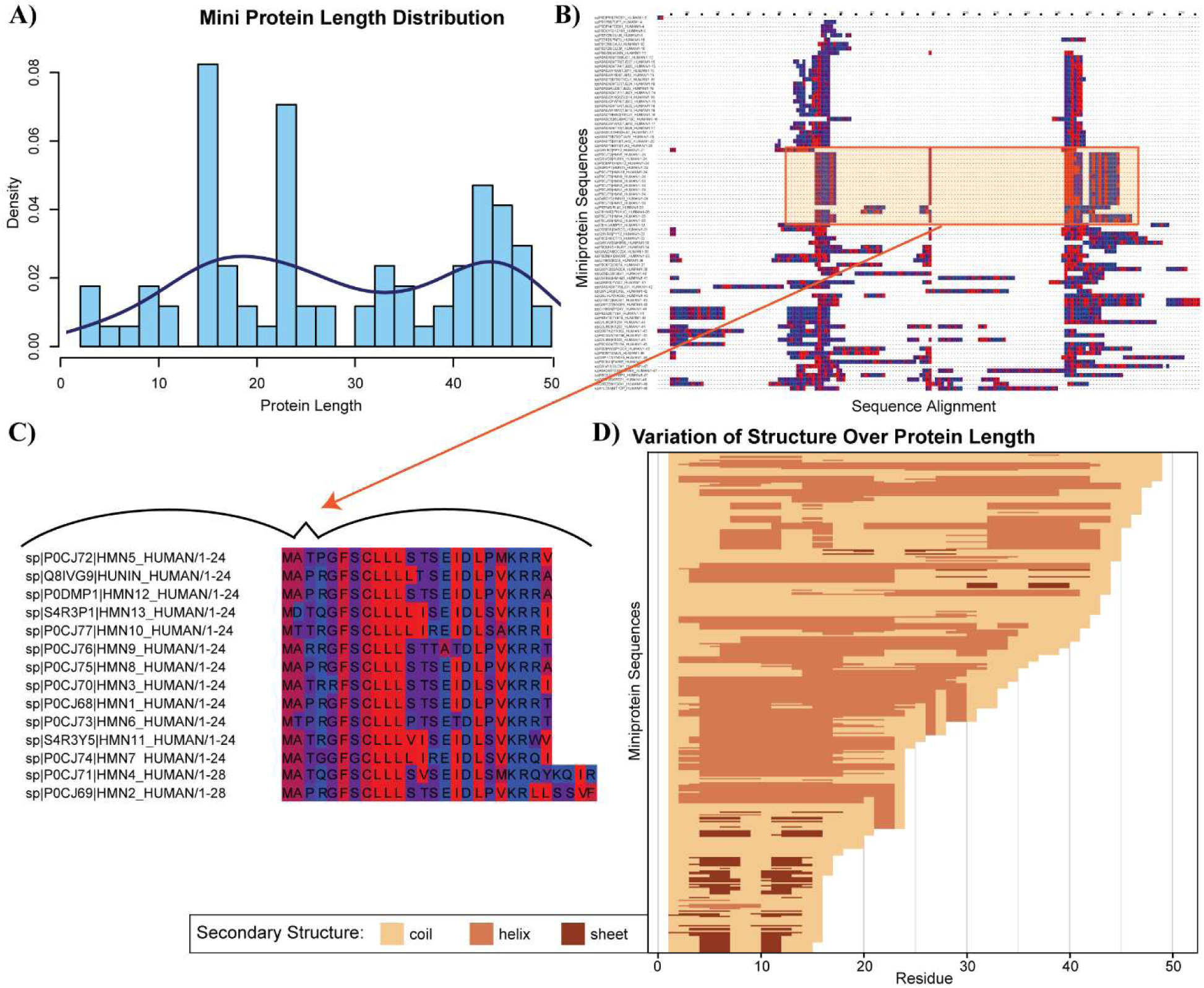
The dataset of miniproteins contains much diversity in length and secondary structure, but miniproteins within a family are often similar length and secondary structure and are well-aligned. A) The length, in number of amino acids, of each miniprotein is shown as a function of density of the broader dataset of miniproteins. B) The sequence alignment of all miniproteins in the dataset is shown as colored by hydrophobicity. C) The humanin and humanin-like family of miniproteins is highly conserved. D) The distribution of secondary structure over each protein sequence is shown. Generally, the longer miniproteins are more helical while the shorter miniproteins mostly contain beta sheets.

**Table 1:** Human Miniprotein Families and Superfamilies from Interpro.

| Families and/or Superfamilies with Miniproteins | # of Human Proteins | Length of Proteins | Miniproteins in Family |
| --- | --- | --- | --- |
| B melanoma antigen-like (BAGE) Family | 5 | 28 | BAGE (1-5) |
| Thymosin Beta Family | 6 | 35-41 | TMSB4 (X & Y), TSMB10, TSMB15 (A-C), |
| Humanin and Humanin-like Family | 14 | 23-24 | MTRNR2L (1-13), MT-RNR2 |
| Keratin-associated protein (KRTAPs) Family | 15 | 41-66 | KRTAP types 8, 19, 20, 21, 22, 23 |
| Metallothionein (MT) | 17 | 25-68 | MT1 (A, B, DP, E, F, G, H, HL1, L, M, X), MT2A, MT (3, 4, B, E, F) |
| Oligosaccharyltransferase Subunit and Complex Proteins | 2 | 37-79 | TMEM258, OST4 |
| Large ribosomal subunit protein eS32 | 1 | 25 | RPL41 |
| Sarcolipin | 1 | 31 | SNL |
| Sarcoplasmic/endoplasmic reticulum calcium ATPase regulator DWORF | 1 | 35 | STRIT1 |
| Short transmembrane mitochondrial protein (STMP) | 1 | 47 | STMP1 |
| Myoregulin (MLN) | 1 | 46 | MRLN |
| Death-like Domain | 19 | 35-76 | *List omitted for length* |
| Immunoglobulin-like Domain (includes T-cell receptor Family) | 104 | 15-79 | *List omitted for length* |

Next, we turn to algorithms designed to identify a different type of protein, namely intrinsically disordered proteins (IDP). Being short polymers, assuming that all miniproteins are IDPs would be natural. However, we anticipate a spectrum from complete IDPs, through meta-stable or metamorphic proteins, to the most stable and ordered protein domains. According to miniprotein gene annotations, nine miniproteins are fully IDP, two contain intrinsically disordered regions (IDRs), and 37 are primarily structured or are not predicted to be disordered. Further, which of these three groups a miniprotein falls into is not strictly length dependent, though all the beta-thymosin family proteins were predicted to be disordered. Therefore, we successfully annotated this diverse set of miniproteins with functional information that indicates groups of related molecules (**Table 2**).

**Table 2:** Selected Miniprotein Family Observations.

| Family or Superfamily | Observations and Notes |
| --- | --- |
| B melanoma antigen-like (BAGE) Family | • Alternating between hydrophobic and neutral (center is hydrophilic) |
| Thymosin Beta Family | • Mainly hydrophilic with hydrophobic residues here and there |
| Humanin and Humanin-like | • MA_R (1-4) --> partially hydrophobic residues to strongly hydrophilic residue<br>• RGFSC (5-9) followed by<br>• Chain of ~4 Leu--> highly hydrophilic<br>• KRR (21-23) --> basic + hydrophilic residues at the end |
| Sarcoplasmic/endoplasmic reticulum calcium ATPase regulator DWORF | • Highly hydrophobic |
| Immunoglobulin-like Domain | • Aromatic residues (5-6) of W or F (F for T-cell receptors)<br>• G_GT / G_GS (7-10) --> hydroxyl group at end<br>• R (11) --> basic residue followed by...<br>• LSV / LTV (13-15) --> hydroxyl group and branched residues<br>• Differences between beta / delta / alpha TCRs |

### Sequence Analysis and Post-Translational Modifications Reveal Functional Insights into Miniprotein Families

Given our initial results supporting miniprotein groupings based on a combination of sequence length and functional annotations, we next aligned all sequences to investigate the sequence-based, structural, and functional relationships amongst these groups and families. The sequence alignment was ordered by length (**Fig. 1B**) and colored by hydrophobicity to determine possible folds for the miniproteins. Multiple groups of similar miniproteins appeared, such as the humanin family of miniproteins, with residue lengths ranging between 24 and 28 amino acids (**Fig. 1C**). Like other groups in the miniprotein dataset, several conserved residues are consistent within each group, and there is diversity in length.

Next, we investigated miniproteins’ post-translational modifications (PTMs) to gain insights into their functional roles and regulatory mechanisms. Of the 85 miniproteins analyzed, 13 were identified to possess previously characterized post-translational modifications (**Fig. 2A**). Our investigation encompassed various miniproteins, including three representatives from the Humanin family—MTRNR2L3, MT-RNR2, and MTRNR2LZ. Additionally, we observed members of the thymosin domain family, which included TMSB10, TMSB15A, TMSB15B, TMSB4X, and TMSB4Y. Noteworthy domain-containing proteins comprised those harboring the sarcolipin domain in SLN and the DUF4435 domain in STMP1. The list also includes HMHB1, KRTAP20-3, and PNAS-138. Phosphorylation emerged as the most prevalent post-translational modification among the miniproteins, with a notable exception being STMP1, wherein succinylation and lysine 35 modifications within the DUF4435 domain were identified. Thus, miniproteins have significant PTM-based regulation.

**Figure 2:**
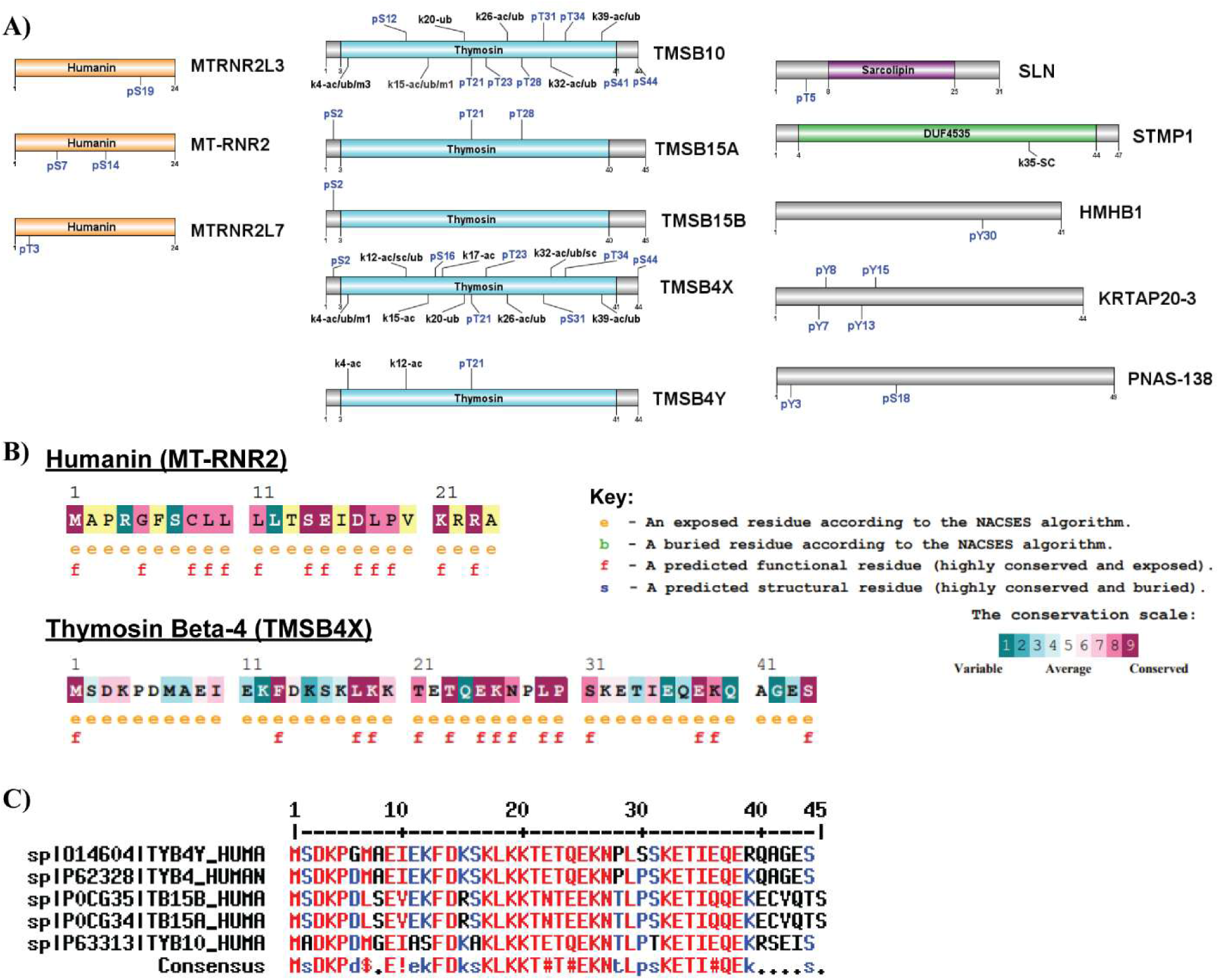
Miniprotein families with a shared domain have many highly conserved residues. A) A domain map for 13 of the miniproteins is shown, with known PTMs and domains noted. B) The conservation and residue properties for one humanin and thymosin-beta protein are annotated. C) An alignment of the thymosin-beta family reveals many conserved residues, especially at the same location as PTMs and functional residues. Red text signifies >90% consensus at that residue; blue signifies >50% consensus.

Focusing on the humanin proteins, we delineated four instances of phosphorylation across the three protein families that exhibited PTMs. Specifically, we identified phosphoserine at position 19 in MTRNR2L3, phosphoserine at positions 7 and 14 in MT-RNR2, and phosphothreonine at position 3 in MTRN2L7. Transitioning to the thymosin proteins, conservation of two phosphorylation sites - phosphoserine 2 and phosphothreonine 21 - was observed. Among these, phosphoserine 2 was detected in TMSB4X, TMSB15B, and TMSB15A, while phosphothreonine 21 was identified in TMSB10, TMSB15A, TMSB4X, and TMSB4Y. Notably, the lysine residue at position 4 was subject to diverse modifications—ubiquitinylation, methylation, and acetylation—in TMSB4Y, TMSB4X, and TMSB10. Further, TMSB4Y and TMSB4X exhibited shared acetylation at lysine 12, while TMSB10 presented a phosphoserine at the same position. TMSB4X and TMSB10 also significantly overlap in lysine modifications at positions 15, 20, 26, 32, and 39, suggesting potential functional synergy between these two miniproteins. Intriguingly, the sarcolipin-containing protein SLN has a phosphorylation at threonine 5, outside the sarcolipin domain.

Since the enzymes that accomplish PTMs bind to their targets in three dimensions, we conducted an evolutionary conservation analysis on the sequences of MT-RNR2 and TMSB4X, representing the humanin and thymosin families of miniproteins, respectively, due to their recurrent PTMs. Within the MT-RNR2 miniprotein, we identified 13 highly conserved residues (**Fig. 2B**). Notably, the serine at position 14 exhibited remarkable conservation, while the remaining PTM sites lacked sufficient data for analysis. This suggests that serine at position 14 holds pivotal functional significance within the humanin domain, with phosphorylation at this site potentially playing a critical role in regulating miniproteins or proteins containing the humanin domain. In contrast, most residues within the thymosin proteins yielded sufficient data for analysis. Residues K4, K20, T21, T23, K26, S31, K32, K39, and S44 exhibited high conservation and were frequently modified (**Fig. 2B** and **2C**). T24, K26, and S44 were the most conserved and modified residues. This suggests that many of these modified residues likely play crucial roles in miniprotein/domain structure and function, emphasizing the importance of T21 and S44, which displayed high conservation and phosphorylation. Furthermore, a multiple sequence alignment of the thymosin proteins revealed that these conserved and commonly modified residues clustered within regions of high consensus, spanning from positions 17 to 27 and 32 to 38. Additional investigations into the remaining modified residues, namely S2, K12, K15, S16, K17, and T34, revealed significant variability. Specifically, S2 was conserved in 4 out of 5 thymosin proteins, except TMSB10, which contained alanine at this position. Further, the lysine at position 12 was observed in 4 out of 5 proteins, with TMSB10 once again being the exception, featuring a phosphorylated serine at this position. Similarly, only 3 of the thymosin proteins exhibited K15, while TMSB15A and B featured an arginine without any modifications at this position. Interestingly, TMSB10 was the sole protein lacking serine at position 16. Finally, positions 17 and 34 were conserved across all thymosin proteins, featuring lysine and thymine residues, respectively. Collectively, this data emphasizes the importance of two modifiable regions in regulating miniproteins or domain structures. Specifically, we highlight phosphorylation at position 21 as a potential crucial regulator of function or structure in these proteins. The variability observed in other sites may significantly confer unique functions to these proteins, such as the phosphorylation of serine 12 in TMSB10 or the absence of lysine 15 in TMSB15 proteins.

### Analysis of Miniprotein Interactome, Pathways, and Networks Reveals Novel Functions and Cellular Roles of Miniproteins

Next, we investigated the potential for miniproteins to act within transient or obligate protein complexes using information derived from 5 high-throughput protein interaction databases. We constructed an interactome for 18 of the 85 miniproteins, with the remainder having no interactions identified. Our analysis revealed 377 experimentally derived or predicted interactions for these miniproteins, consisting of a diverse range of 24 to 49 residues **(Fig. 3A).** A network analysis of the miniprotein interactomes demonstrated that they were largely independent and hence may share few direct targets. However, we observed that the TMSB proteins, specifically TMSB4Y, TMSB10, and TMSB4X, shared a total of 9 interactions, with TERF1, TERF2IP, and POT1 being shared targets among these three miniproteins.

**Figure 3:**
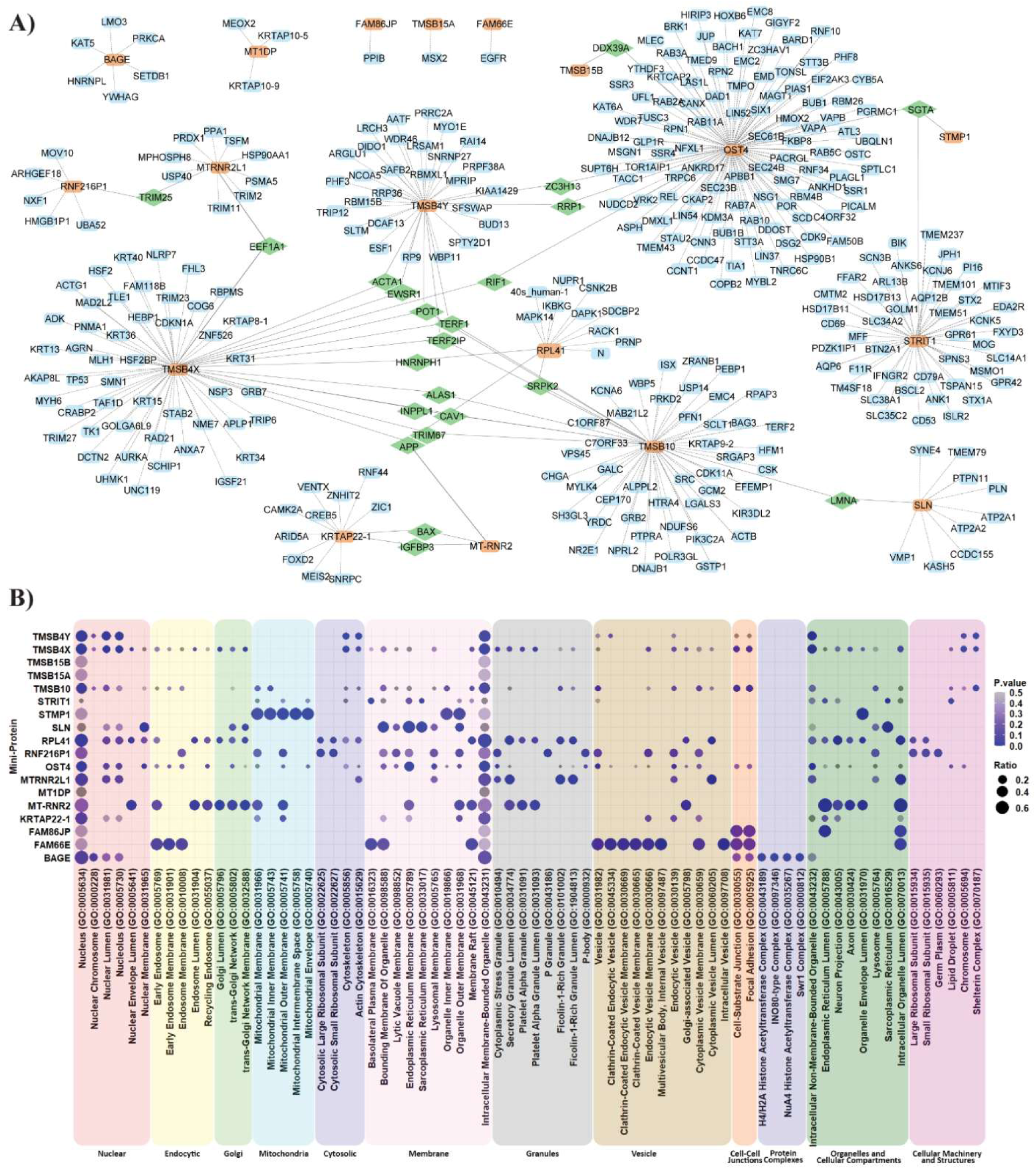
Interaction Network and Gene Ontology Enrichment Analyses reveal connections between miniproteins, suggesting shared functions and/or pathways. A) The interactions that all miniproteins in this study are known to have are shown, with the study’s miniproteins in orange, interactors in blue, and interactors that connect two miniproteins in green. Dashed lines show the interactions themselves. B) The proteins that interact with each miniprotein underwent gene ontology enrichment analysis, and the cellular compartment associated with the miniprotein interactors is shown. Larger circles demonstrate that more of the interactors associate with that compartment; the circle’s color corresponds to the p-value of enrichment.

We performed a gene ontological analysis focusing on the cellular component to gain insights into the functional implications of these miniproteins based on their interactions (**Fig. 3B**). The study revealed distinct enrichment patterns for different miniproteins. The Thymosin (TMS) family of proteins is involved in preventing actin polymerization but has been recently linked to tumorigenesis and progression and thus may play a much more significant role than initially described (Xiong et al. 2022). This family of proteins is enriched in terms related to the nucleus and intracellular membrane-bounded organelle, with TMSB15B and TMSB15A as the only two members that did not reach the significance cut-off of 0.05. Additionally, TMSB10, TMSB4X, and TMSB4Y all showed significant enrichment for the shelterin complex, which consists of 6 telomere-specific proteins and is significantly enriched through the shared interactions with TERF1, TERF2IP, and POT1. Considering that TERF1 and POT1 are a part of the shelterin complex, these three thymosin miniproteins likely play a substernal role in regulating this complex, protecting telomeres from DNA repair mechanisms, and regulating telomerase activity. Thus, here we show that the TMSB proteins are located within the nuclear component and may act on the shelterin complex, and this dysregulation of the TMSB proteins may disrupt the shelterin complex, resulting in the deterioration of telomere integrity.

The Humanin protein MT-RNR2 has been observed with a variety of functions, from suppressing apoptosis, inhibiting fibril formation, reducing inflammation, aiding mitochondria biogenesis, and enhancing insulin sensitivity (Alsanousi et al. 2016; Guo et al. 2003; Muzumdar et al. 2009; Sreekumar et al. 2016; ZhaoZhao & Li 2013). Thus, it was surprising that the interactome of MT-RNR2 exhibited enrichments for an extensive range of terms within the Golgi and lumen-associated groups. This may suggest indirect interactions or signaling pathways involving Golgi or cellular lumen structures concerning Humanin’s functions. Similarly, the human-like 1 protein MTRNR2L1 showed enrichments for lumen-based terms like cytoplasmic vesicle lumen and intracellular organelle lumen as well as granular lumen terms such as secretory granule lumen, ficolin-1-rich granule, and filling-1-rich granule lumen, further highlighting a potential mechanism between humanin and humanin-like proteins with or cellular lumen structures.

As for the sarcolipin-related proteins, STRIT1 (Small transmembrane regulator of ion transport 1) is a P-type ion transporter responsible for sequestering cytoplasmic Ca^2+^ into the sarcoplasmic reticulum (SR) of cardiac muscle cells (Cleary et al. 2022). STRIT1 showed significant enrichment for the basolateral plasma membrane and sarcoplasmic reticulum membrane, with the latter being expected due to the known function of the protein. Similarly, Sarcolipin (SLN), previously shown to regulate Ca-ATPase, showed high expected enrichment terms in the sarcoplasmic reticulum, specifically the sarcoplasmic reticulum membrane (Mascioni et al. 2002). Other terms were related to various cellular membranes of different components, such as the bounding membrane of organelles, endoplasmic reticulum membranes, organelle outer membrane, and nuclear membrane. This suggests that sarcolipin and STRIT1 may have roles in other membranes or that these annotations are picking up on their general membrane-related features.

Little is known about proteins Sequence Similarity 86 Member A pseudogene (FAM86JP) and Sequence Similarity 66 Member E (FAM66E). Both showed enrichments for the cellular component cell-substrate junction and focal adhesion; additionally, FAM66E was observed to have enrichments within the vesicle grouping, specifically for the terms vesicle, clathrin-coated endocytic vesicle, clathrin-coated endocytic vesicle membrane, endocytic vesicle membrane, multivesicular body internal vesicle, cytoplasmic vesicle membrane, and intracellular vesicle. With no prior studies describing the functions of FAM86JP or FAM66E, we infer that these two miniproteins may be involved in endocytic processes and intracellular vesicle dynamics based on their comprehensive interactome.

The Short Transmembrane Mitochondrial Protein 1 (STMP1) is a mitochondrial peptide observed to enhance mitochondrial fission and thus promote tumor cell migration (Xie et al. 2022). STMP1 displayed significant enrichment associated with mitochondria, including mitochondrial membrane, inner membrane, outer membrane, intermembrane space, and mitochondrial envelope. Given its known functions, this is expected and demonstrates that our network analysis can highlight actual functionality.

Our analysis also revealed that the ribosomal large subunit protein RPL41 exhibited expected enrichment within cytosolic groups such as large and small ribosomal subunits. However, the most significant enrichment was observed within the granules group, particularly in the secretory granule lumen and ficolin-1-rich granule. Furthermore, the Putative protein RNF216-like (RNF216P1) displayed similar enrichments within the granule group but also featured terms associated with cytoplasmic stress granules, P-granules, and P-bodies, along with the same cytosolic terms. These unexpected findings reveal a possible connection between the well-studied RPL41 and less-understood RNF216P1 miniproteins.

Dolichyl-diphosphooligosaccharide--protein glycosyltransferase subunit 4 (OST4) is a subunit of the mammalian oligosaccharyltransferase required for efficient N-glycosylation has been previously identified to be more than 50% embedded into the endoplasmic reticulum membrane; thus, enrichments for endoplasmic reticulum and intracellular membrane-bounded organelles is expected (Dumax-VorzetRoboti & High 2013). However, beyond the expected findings, we also noted the enrichment of some of the OST4 interactors involved with the nuclear cellular component, suggesting that these proteins may be modified post-translationally by OST4 after protein synthesis in the endoplasmic reticulum.

Lastly, B melanoma antigen 1 (BAGE1) was generated by juxta-centromeric reshuffling of the KMT2C/MLL3 gene; while its function remains unknown, it is an antigen found in human melanomas (Ruault et al. 2003). Cellular component ontological analysis derived from the comprehensive interactome established in this study displayed enrichments for nuclear proteins, specifically HNRNPL, SETDB1, KAT5, and LMO3. KAT5 was also found enriched for protein complexes the H4/H2A histone acetyltransferase, INO80-Type, NUA4 histone acetyltransferase, and Swr1 complexes, suggesting a potential involvement of BAGE1 in chromatin remodeling and histone acetylation processes mediated by BAGE1’s interaction with KAT5. MT1DP and KRTAP22-1, however, did not exhibit significant enrichment in the cellular components analysis. While our analysis of miniprotein interactors and their respective associated terms revealed many expected findings, novel connections were also found, which may further explain the function of less studied miniproteins. This methodology could be helpful in further analysis of miniproteins.

### Peptide-Based Algorithms May Outperform Deep Learning Methods for Miniproteins

The cutoff between when a short amino acid polymer behaves like a peptide versus a developing secondary or tertiary structure is unclear and likely not strictly determined by polymer length. Thus, we next sought to characterize the relative performance in structure prediction for peptide-based algorithms compared to the new AI-based structure prediction algorithms, using PEPFOLD3 and AlphaFold2, respectively. Two metrics, TM and Qmean scores, were used to compare protein structures generated by the two structure prediction tools. TM scores (Zhang & Skolnick 2004), which quantify overall topological similarities, were compared (**Fig. 4A**). PEPFOLD3 had relatively better TM scores than AlphaFold2 for most miniproteins (∼90%). The second metric used to compare the two tools was Qmean (BenkertTosatto & Schomburg 2008), which quantifies the geometrical aspects of protein structures (**Fig. 4B).** In contrast to TM scores, Qmean scores were relatively similar between the two algorithms and evenly distributed across the range of miniprotein sizes, with two outliers being 25 amino acids long. We next assessed the size associations with structure quality in more detail. Regarding TM scores, miniproteins of the smallest size seem to have worse scores in AlphaFold2 than in PEPFOLD3. PEPFOLD3 Qmean scores also supported this finding, with smaller miniproteins having comparatively better scores than larger miniproteins. AlphaFold2 Qmean scores had no discernable association with miniprotein size. Ultimately, this analysis implies that protein size has a role in selecting structure prediction algorithms. Consequently, miniprotein analysis and peptide-based and AI-based approaches can be used together to inform about structural features. Still, peptide-based approaches will likely be more accurate for miniproteins, especially smaller ones.

**Figure 4:**
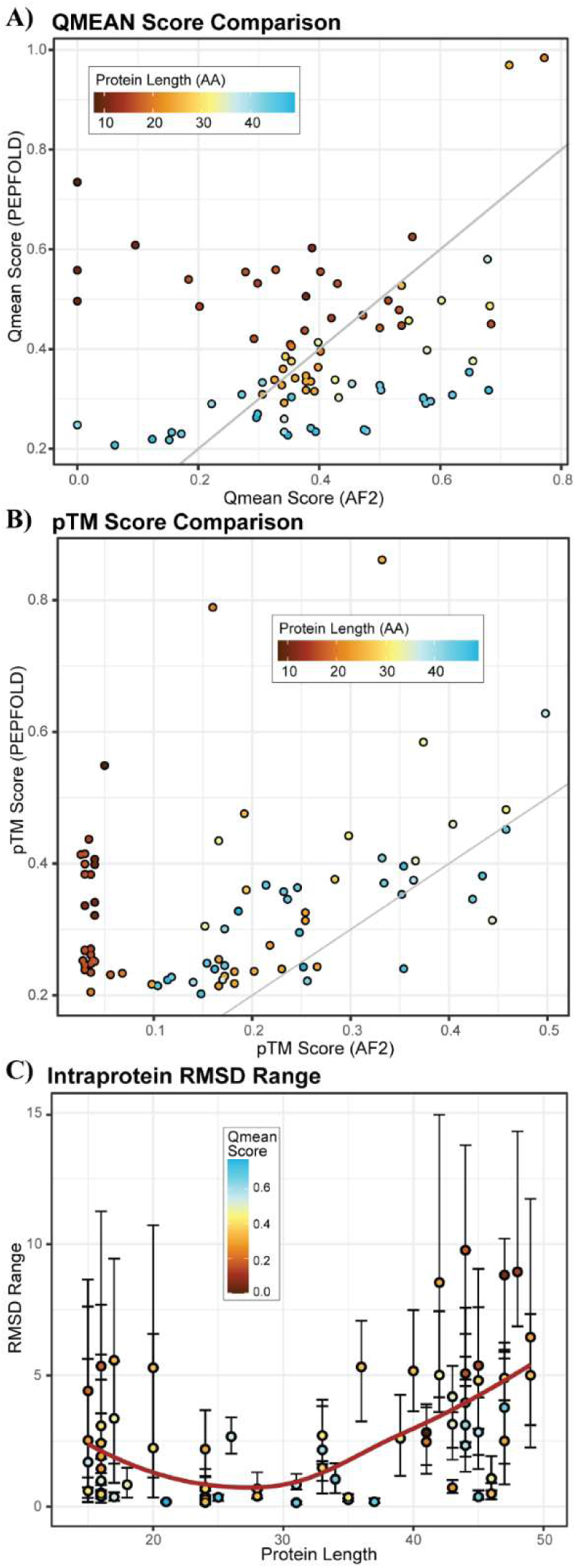
Shorter miniproteins have more diverse structures and are better predicted by PEPFOLD than Alphafold2. A) the QMEAN score of PEPFOLD and AlphaFold2 for each predicted protein structure are plotted against one another, with each protein colored by its length. B) The TM score of PEPFOLD and AlphaFold2 for each predicted protein structure are plotted against one another, with each protein colored by its length. C) The RMSD between the PEPFOLD and AlphaFold2 models of each protein is plotted as a function of that protein’s amino acid length, and each point is colored by the QMEAN score.

It was found that miniproteins on both ends of the length scale differed more in structure alignment, with an overall higher root mean square deviations (RMSDs) and range of scores across the multiple models generated by each approach. Factoring in how quality score affects these results, we could see that more confident scores, according to each tool, generally resulted in more consistent structures (**Fig. 4C**). Those on the lower and higher end of the size scale had more structural diversity across predicted models. Even between miniproteins, which are very closely related, we find vast differences across their structures. For example, humanin miniproteins have similar sequences but their structures are divergent (**Fig. 5B**) and sometimes have high RMSDs compared to each other (**Fig. 5A)**. The same pattern exists for the T-cell receptor proteins (**Fig. 5C-D**), where there is evident diversity despite the similarity of the groups. These differences in protein sequence and structure may imply different or diverging functions. Further study of the differences between related miniproteins and the causes thereof may be enlightening on miniprotein functions, diversity, and evolution.

**Figure 5:**
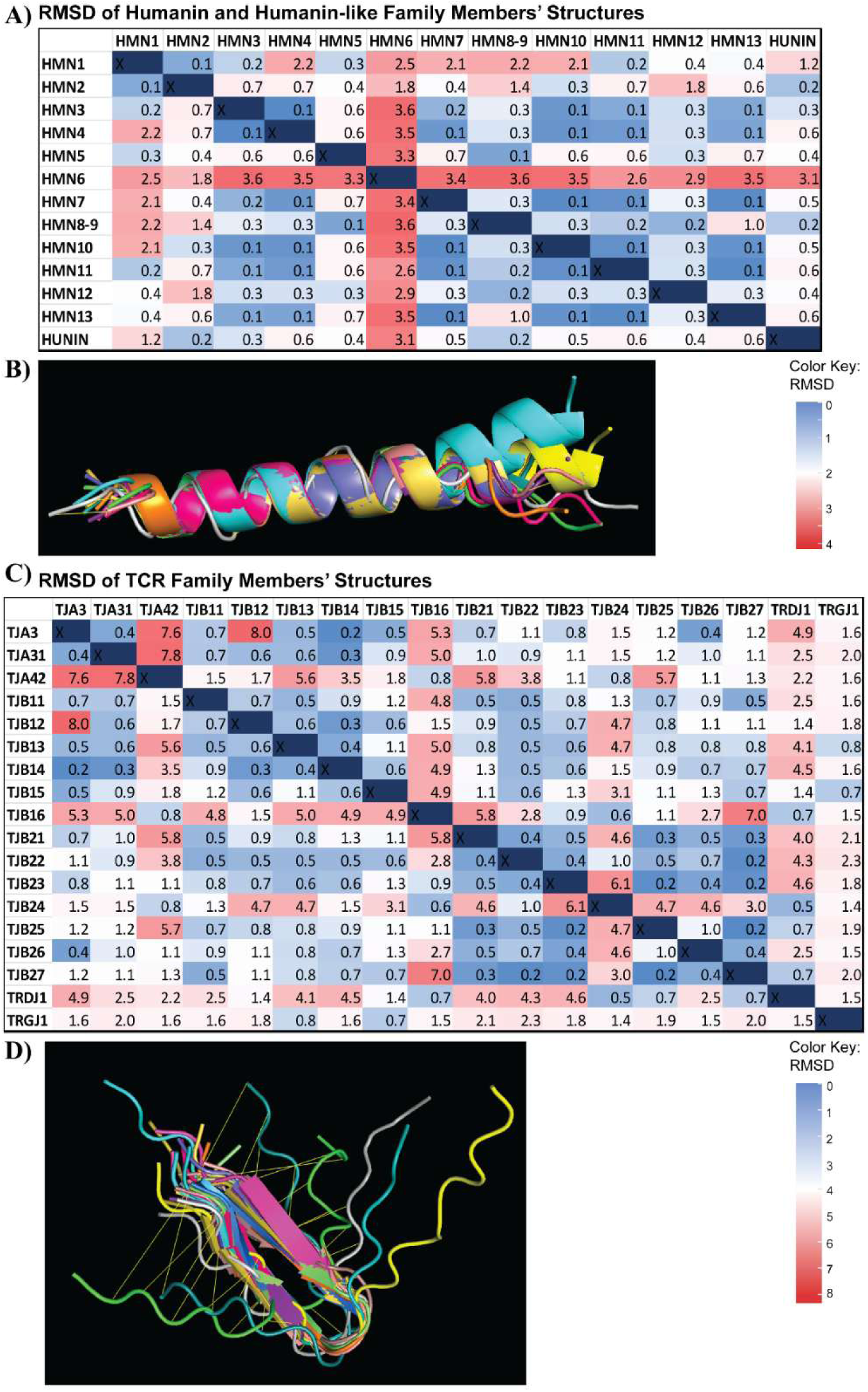
Miniproteins within the same family have varying degrees of structural similarity despite having small structures overall. A) the RMSD of each humanin and humanin-like family protein against all other proteins in the family is shown. B) all humanin protein structures are aligned and are visually quite similar. C) the RMSD of each T-cell receptor (TCR) family protein against all other proteins in the family is shown. D) all TCR protein structures are aligned and some structures are visually quite distinct.

### Varying Structural Motifs and Structure Diversity Amongst Miniproteins

Next, we assessed secondary structure variability. Visible secondary structures were recorded for each miniprotein and its corresponding top 5 structural models. To identify possible structure trends within the miniprotein dataset, structural motifs for each miniprotein were plotted by size (**Fig. 1D**). Hairpin loop structures, also known as beta-turn-beta motifs, were common amongst smaller miniproteins. There was also a considerable presence of alpha-helical structures, seen in the more medium-to-large miniproteins. More well-known protein motifs, such as the helix-turn-helix motif, were also observed and were representative of DNA-binding proteins. Additionally, some secondary structures represented miniprotein families, which were identified in our preliminary findings. For example, the T-Cell Receptor miniproteins (15-20 aa) typically contained hairpin loop structures, and the Humanin protein family (24-28 aa) all contained alpha-helical structures. However, we also found that 24% of the miniprotein structures were entirely coiled, with no regular three-dimensional structure, including at least one structure of all proteins predicted to be disordered, excluding the thymosin-beta family (which all had alpha helices). Thus, both peptide-based and AI-based structure prediction algorithms should play a role in functionally annotating miniproteins.

### Structural Similarity Between Miniproteins and Larger Protein Domains Reveals Potential Functional Mimicry

Finally, we compared structures between these predicted miniprotein structures and solved protein structures in the Protein Data Bank (PDB) to assess how well the modeled structures resemble experimentally determined folds. This allowed us to identify potential structural alignments and investigate the overlap of functional residues. Due to a lack of length (the software requires at least 30 amino acids in the structure) and other restrictions, 19 of the 85 miniproteins returned results. The results consisted of 67 unique protein structures that have a shape-based resemblance to a miniprotein predicted structure. The RMSD scores and Z-scores between the miniprotein and more extended protein structures were recorded and put onto scatterplots, showing progression based on miniprotein length (**Fig. 6A-C).** While there aren’t any strong correlations between size and structure prediction scores, we found several structures that were highly similar to structures in our miniprotein dataset. For example, Uncharacterized Protein PRO1716 has a z-score of 5.4 when matched with a motor protein structure (MYH7 residues 1526-1571 fused to Gp7, a chimera protein) (**Fig. 6D**). On the other hand, HR upstream open reading frame protein (HRURF) had a low Z-score of 2.2 when structurally aligned to the recombination protein MRE11 in complex with RAD50 and ATP (**Fig. 6E)**. Though these Z-scores are on opposite ends of the spectrum, both incorporate almost the entire miniprotein with their matched structure, thus providing additional support for the feasibility of the predicted structures and next steps for future research to interrogate the likely function of these miniproteins.

**Figure 6:**
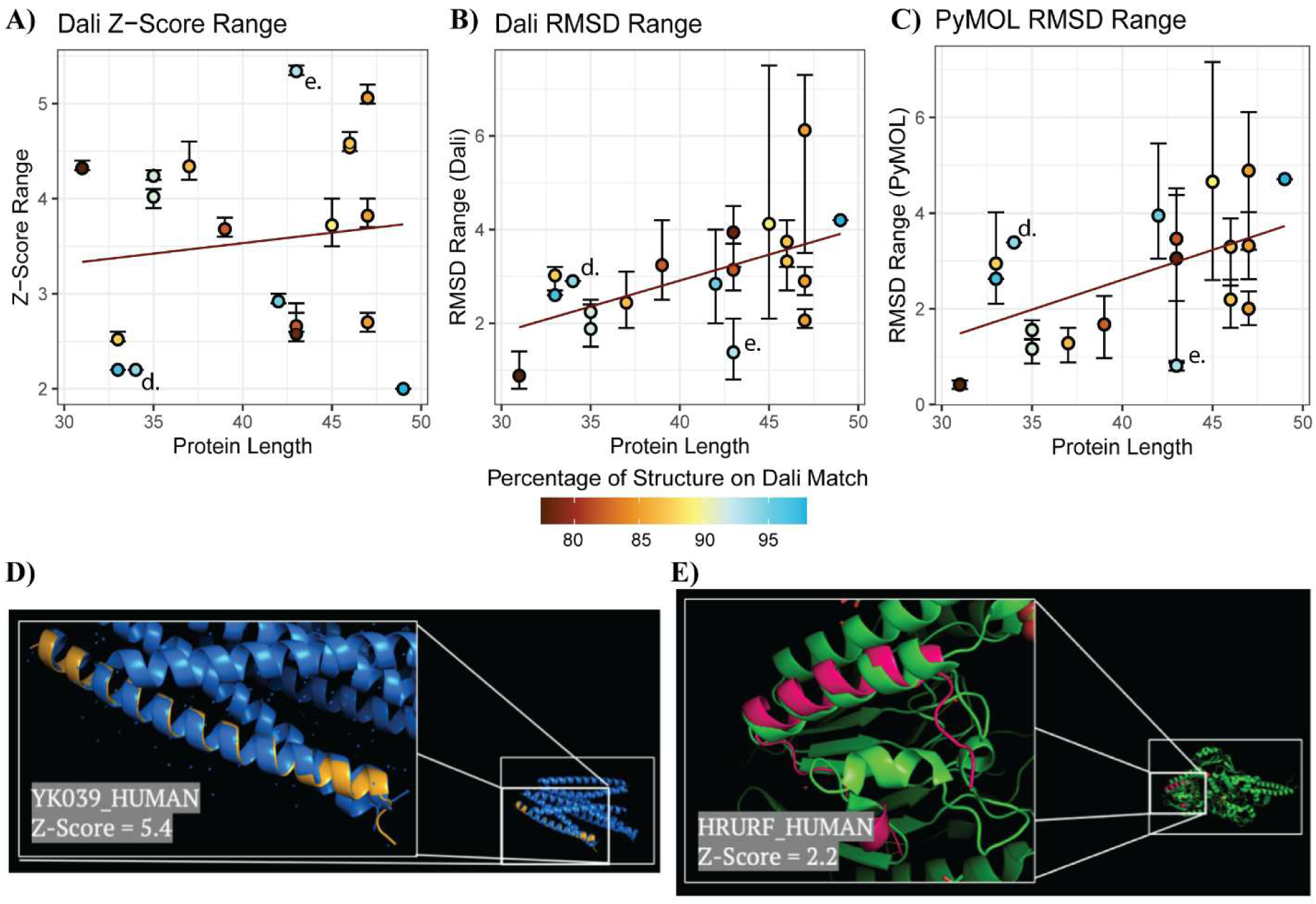
Miniprotein structures are similar to other protein structures. A) The z-score of Dali’s most similar structure to each miniprotein is shown, colored by the percent of the miniprotein structure matched to the other structure. B) The RMSD, as calculated by Dali, of Dali’s most similar structure to each miniprotein is shown, colored by the percent of the miniprotein structure matched to the other structure. C) The RMSD, as calculated by PyMOL, of Dali’s most similar structure to each miniprotein is shown, colored by the percent of the miniprotein structure matched to the other structure. D) The structure of miniprotein PRO1716 is shown aligned to a MYH7 structure Dali found. The position of this match is marked by “d.” on A, B, and C. E) The structure of miniprotein HRURF is shown aligned to a MRE11 structure Dali found. The position of this match is marked by “e.” on A, B, and C.

## Discussion

This study makes notable contributions to the field of genomics by analyzing human miniprotein functional diversity, which additionally clarifies that they are not dichotomous (i.e., structured versus unstructured), and their structure propensity is not strictly dependent on length. Additionally, the context specificity of miniprotein three-dimensional structure may be more robust than for globular proteins due to their sequence composition differences. These differences in miniproteins may indicate evolutionary occurrences, thus possible differences in their structure and function. Structural scoring and analysis were performed to support these conclusions.

The revolution of AI-based algorithms for protein structure prediction has driven innovations, yet the limits of these algorithms need to be understood better. We found that peptide-based algorithms may outperform deep learning methods in predicting miniprotein structure. Additionally, the predicted forms of miniproteins mimic topologies observed in larger molecules. This supports the feasibility of the predicted models and suggests that miniproteins could play a competitive regulatory role in larger complexes. However, in many cases, the identity and function of those larger proteins are still unknown and must be further studied.

Through this utilization of several sequence annotation tools, we also observed the difference in annotations between web tools dedicated to proteins and those dedicated to genes. The lack of protein annotations supports the notion that miniproteins are heavily understudied. Meanwhile, there is a significantly larger number of annotations and scores for the genes that encode the miniproteins–likely from more commonly performed genomic studies. These annotations revealed common traits and high conservation within miniprotein families, yet substantial diversity despite each sequence being so short. Annotating miniproteins can lead to identifying the potential functionality of miniproteins – as we were able to do for 14 miniproteins, proposing novel or additional functions for 10 miniproteins and finding concordant functional data for the other 4. Our findings highlight the diverse and often unexpected functions of miniproteins, expanding our understanding of their biological significance and potential roles in cellular processes. Further, the fact that these previously disregarded proteins seem to play such pivotal roles in critical processes such as chromatin remodeling and telomerase regulation demonstrates the clear significance of miniproteins.

## Conclusions

In conclusion, we found many miniproteins fold into common secondary structure motifs. We also found a wide variety of interactions and likely functions across the miniprotein in our cohort, revealing a greater diversity and importance than is sometimes attributed to these short proteins. This work has theoretical implications on how we conceptualize and design miniproteins, and practical implications for selecting the best tools to study miniproteins. Miniproteins are important to understand so we can identify new contributors to physiology, pathobiology, and diversity advanced drug design. Therefore, we anticipate that the field of genomics will increasingly study additional dimensions of miniprotein function.

